# Pro-social and pro-cognitive effects of LIT-001, a novel oxytocin receptor agonist in a neurodevelopmental model of schizophrenia

**DOI:** 10.1101/2023.05.24.542076

**Authors:** Diana Piotrowska, Agnieszka Potasiewicz, Piotr Popik, Agnieszka Nikiforuk

## Abstract

Social and cognitive dysfunctions are the most persistent symptoms of schizophrenia. Since oxytocin (OXT) is known to play a role in social functions and modulates cognitive processes, we investigated the effects of a novel, nonpeptide, selective OXT receptor agonist, LIT-001, in a neurodevelopmental model of schizophrenia. Administration of methylazoxymethanol acetate (MAM; 22 mg/kg) on the 17^th^ day of pregnancy is known to cause developmental disturbances of the brain, which lead to schizophrenia-like symptomatology in the offspring. Here, we examined the effects of acutely administered LIT-001 (1, 3, and 10 mg/kg) in MAM-exposed males and females on social behaviour, communication and cognition.

We report that MAM-treated adult male and female rats displayed reduced social behaviour, ultrasonic communication and novel object recognition test performance. LIT-001 partially reversed these deficits, increasing the total social interaction time and the number of ‘happy’ 50 kHz ultrasonic calls in male rats. The compound ameliorated MAM-induced deficits in object discrimination in both sexes.

Present results confirm the pro-social activity of LIT-001 and demonstrate its pro-cognitive effects following acute administration.

## INTRODUCTION

Although the ‘positive’ symptoms are the most characteristic aspect of schizophrenia, the ‘negative’ and cognitive ones are the most difficult to address and are persistent even in the periods of remission of psychosis. Negative symptoms are usually described as depressive-like but also include social withdrawal and difficulties in verbal communication (Messinger et al., 2011). Antipsychotics with the typical (antagonism of mesolimbic dopamine D2 receptors) and atypical (antagonism of D2 and serotonin 5- HT2A receptors) mode of action have little effect on the negative symptoms and cognitive deficits. Thus, there is a need for new therapeutics addressing the social and cognitive domains of schizophrenia symptoms (Millan et al., 2016). New ideas come every decade, and recently, there has been a focus on oxytocin’s system involvement in the aetiology of some schizophrenia-related deficits (for review, see Shilling & Feifel, 2016).

Oxytocin (OXT) was first described by Sir Henry Dale in 1906, and its nonapeptide structure was discovered by Du Vigneaud (1953). OXT is most known for its crucial role in inducing birth and lactation, but it also works as a neuromodulator in the brain and has been reported to have effects on numerous functions, mainly social recognition and memory (human studies: Guastella et al., 2009; Rimmele et al., 2009; Savaskan et al., 2008; animal studies: Ferguson et al., 2001; Popik & van Ree, 1991), as well as spatial memory (Maier at al., 2016), stress reduction and coping (e.g., in PTSD: Olff et al., 2007), anxiety reduction (reviewed by Amico et al., 2004), regulation of aggressive behaviour (DeVries et al., 1997; Gulevich et al., 2019) and inflammatory pain (Hilfiger et al., 2020). The current state of knowledge on oxytocin’s function in the brain has been recently reviewed by Grinevich and Neumann (2020).

Numerous attempts have been made to use OXT to modulate social functioning in patients with social phobia, autism spectrum disorders, and schizophrenia. The history of oxytocin in schizophrenia research has been extensively reviewed by Feifel et al. (2016). Since the peptide structure of oxytocin is well known, the easiest way to use it in the clinic is to administer synthetic oxytocin or its analogue, but this method has some limitations. The first is OXT’s moderate brain permeability and short half-life in the tissue (Gossen et al., 2012; Ring et al., 2010; for review, see Guastella et al., 2013). OXT in nasal spray is commercially available despite the lack of scientific consensus on its beneficial effects. Another limitation is the evolutionary similarity of oxytocin and vasopressin systems. Although there is only one known OXT receptor, OXT’s affinity to vasopressin receptors (VRs) must be taken into consideration (Busnelli & Chini, 2018). The precise effects of OXTR functions are difficult to distinguish from the effects of VRs’ activation due to the lack of selective OXTR agonists (for review, see Dumais & Veenema, 2016). Hence, there is a need for a selective, nonpeptide OXTR agonist. Such an agonist, LIT-001, was synthesised by Frantz et al. (2018) and tested in a mice model of autism spectrum disorder with promising results.

Schizophrenia is currently described as a heterogenous neurodevelopmental disorder with a strong genetic component, suggesting that its core relies on alterations in the brain resulting from disturbances occurring during the period of brain development (first proposed by Weinberger in 1986, revised and reviewed by Weinberger (2017). An approach addressing this aspect in preclinical research is to disturb the development of the rat brain at an early stage and observe the schizophrenia-like symptoms emerging later in life. One of the neurodevelopmental models relies on mitotoxin methylazoxymethanol acetate (MAM) administration on gestational day 17, which results in biochemical, neuroanatomical, and behavioural changes (recently reviewed by Maćkowiak, 2021).

One of the most commonly used paradigms to investigate the social competencies of animals is the social interaction test (SI), which allows observation of two freely behaving rats (for review, see Gururajan et al., 2010). It has been reported that in pharmacological and neurodevelopmental schizophrenia models, the time and the pattern of social behaviours were distorted (Flagstad et al., 2004; Potasiewicz et al., 2018). Rats communicate with conspecifics with a variety of different calls within the ultrasonic spectrum (Panksepp & Burgdorf, 2000). Such vocalisations can be recorded and analysed. The SI test also allows for the recording of ultrasonic vocalisations (USVs). The analysis of USVs gives an insight into animals’ emotional state, as the vocalisations reflecting negative and positive affective states are in the different ultrasonic spectrums: the ‘alarm’ calls are of about 22 kHz, while the ‘happy’ calls are produced at the frequencies of ∼ 50 kHz (for review, see Brudzynski, 2015). Within 50 kHz calls, several sub-categories can be distinguished (Wright et al., 2010).

A novel object recognition test (NOR) serves to assess cognitive impairments in animal models. In the NOR test, animals’ ability to recognise a previously presented object is scored by measuring the time spent interacting with a novel and a familiar object. An inability to distinguish a novel from a familiar object is considered a cognitive deficit (Ennaceur & Delacour, 1988; Nikiforuk et al., 2013). It was previously shown that in various rodent schizophrenia models (e.g., genetic, pharmacological, and developmental), the ability to distinguish novel from familiar objects in this test was disrupted (for review, see Lyon et al., 2012).

Previously, we have shown social and cognitive deficits in the MAM model (Potasiewicz et al., 2018), and in another paper, a reduction of oxytocin concentration in the MAM-treated rat brains was reported (Potasiewicz et al., 2020). In the present study, we investigated whether social and cognitive impairments could be alleviated with the novel, nonpeptide, selective OXTR agonist - LIT-001.

## MATERIALS AND METHODS

### Animals

Fourteen pregnant dams (Sprague Dawley rats) were obtained from Charles River (Sulzfeld, Germany) on gestational day (GD) 15 and housed individually in polycarbonate cages (length × width × height: 42 × 26.5 × 18 cm). On GD 17, the dams were intraperitoneally injected with vehicle (VEH, 0.9% NaCl; 7 females) or a single dose of 22 mg/kg MAM (7 females) in a volume of 1 ml/kg. The gestation of MAM-treated dams proceeded normally and terminated on the 21st–22nd day. The total number of born animals was 195. MAM treatment did not affect litter size (10–17 pups) or pup survival (Supplement, Fig. S1). On the postnatal day (PND) 21, all pups were weaned and assigned into same-sex, same-litter groups of 4–5 rats.

Females and males were housed in different rooms under standard laboratory conditions (12-h light/dark cycle from 6 a.m., with a controlled temperature of 21 ± 1 °C and humidity of 40–50%). After weaning, offspring were handled three times a week. All of the behavioural tests were performed during the light phase (9 a.m.–5 p.m.) of the light/dark cycle (Monday to Friday). Animals had *ad libitum* access to food and water.

The experiments were conducted in accordance with EU directive 2010/63/EU and were approved by the Ethics Committee for Animal Experiments, the Maj Institute of Pharmacology, no. 201/2019.

### Drug preparation

Methylazoxymethanol acetate (MRI Global, Kansas, US) was dissolved in NaCl (0.9%) and injected intraperitoneally (i.p.) in a volume of 1 ml/kg to pregnant rat dams.

LIT-001 (MedKoo Biosciences, Morrisville, USA), a very specific agonist for OXTR with limited off-targets and a long-lasting (>2 h) half-life, was prepared as described in Frantz et al. (2018) and Hilfiger et al. (2020). In short, LIT-001 was dissolved in carboxymethyl cellulose (CMC, 1%) - NaCl (0.9%) and administered at the dose of 1, 3, or 10 mg/kg. LIT-001 or vehicle were injected i.p., in a volume of 2 ml/kg, 30 min before each test.

### Experimental Design

To assess social and cognitive functions, male and female rats prenatally treated with either VEH or MAM were subjected to the SI and NOR tests. The experimental design is presented in Fig. 1A. To minimise the risk of a ‘litter effect,’ rats were randomly distributed across experimental groups. Males and females were tested on separate days, males first and females on the following day. To reduce the number of needed animals, the SI and NOR tests were done twice, with a week washout period between tests. For the SI test, animals were assigned to the new pairs, and for the NOR test, a different set of objects was used in the second test. No animal received the same treatment or dose twice.

**Figure 1.**
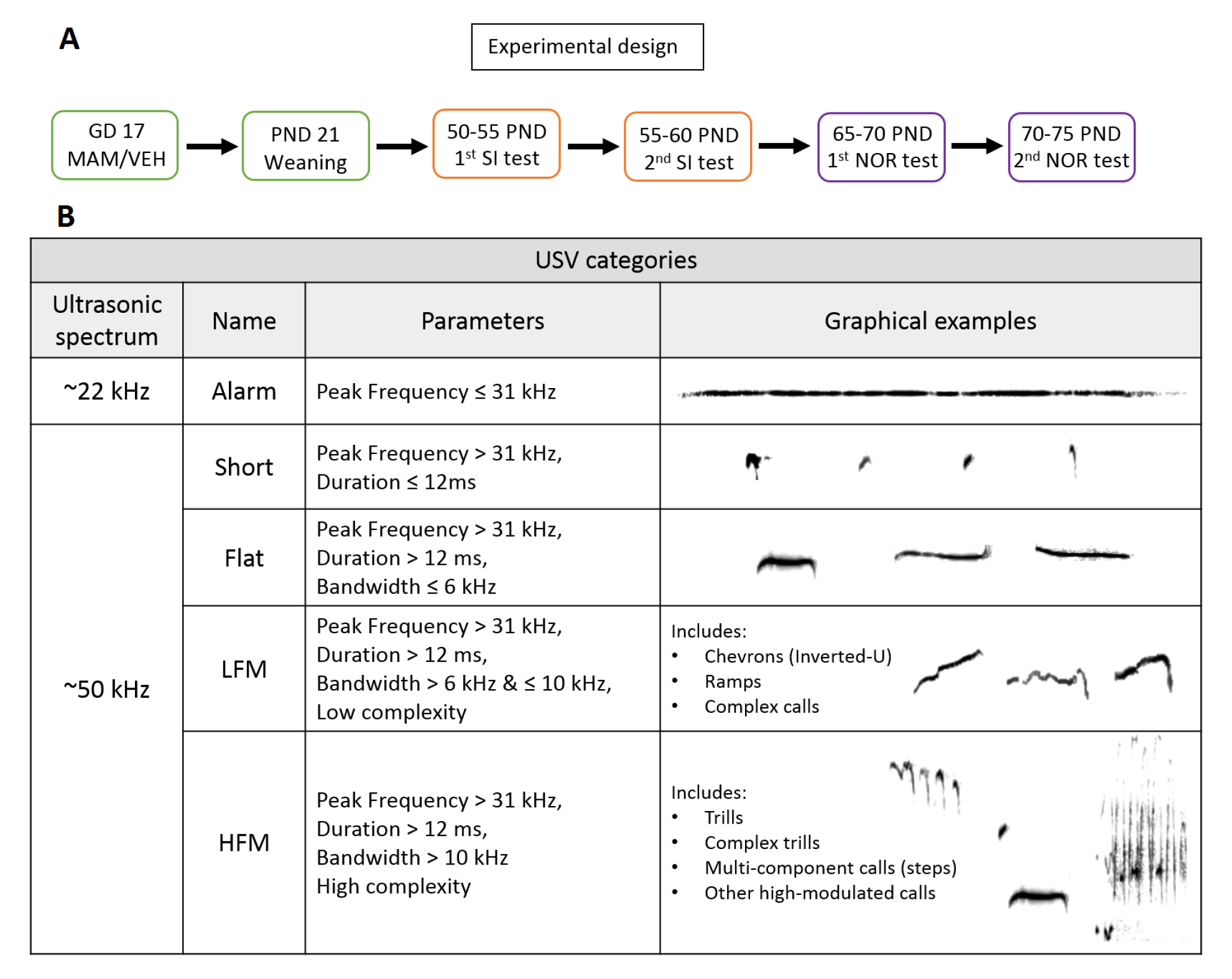
Experimental design (A) and Ultrasonic vocalisations categories (B). ***A**. Graphical explanation of the experimental design. Abbreviations: GD - gestational day, MAM - methylazoxymethanol acetate, NOR - Novel Object Recognition test, PND - postnatal day, SI - Social Interaction test, VEH – vehicle. **B**. Ultrasonic vocalisations categories with graphical representations. Abbreviations: HFM – high frequency modulated, LFM – low frequency modulated, USV – ultrasonic vocalisations. Modified from Potasiewicz et al. (2020).*

One day before the Social Interaction test, animals were habituated to the testing room and apparatus. Adaptation to the testing apparatus (open field arena made of black Plexiglas, length × width × height: 57 × 67 × 30 cm) was also recorded and analysed using the Any-maze® tracking system (Stoelting Co., Illinois, USA) to measure rats’ baseline exploratory behaviour.

### Social Interaction test (50-60 PND)

The Social Interaction test was performed as previously described (Hołuj et al., 2015). In brief, animals were assigned into pairs under the criteria of the same sex, same treatment, matching body weight (+/- 10g), different home cages, and preferably different litter. Each pair was placed in an open field arena (described in details in the section Experimental Design) for 10 min. The behaviour was recorded by a CCTV camera placed above the arena and connected to the Any-maze® tracking system. The arena was thoroughly cleaned after each test.

Social behaviours were scored manually by an experienced observer blinded to the treatments. The following social behaviours were scored: sniffing, anogenital sniffing, grooming, following, and climbing. The time spent in active social behaviours was summed to yield a total score. Because both animals in a pair yielded approximately equal scores, the social interaction time was expressed as a summed score for each pair of animals. All behaviour analysis was done in the Noldus Observer XT, version 10.5 (Noldus Information Technology, Wageningen, The Netherlands). Simultaneously, social interaction-induced vocalisations were recorded by an ultrasound microphone (for details, see the ‘USVs: recording and analysis’ section).

### USVs: recording and analysis

For the recordings of the vocalisations, we used an ultrasound microphone with a frequency response range of 2 kHz – 200 kHz (UltraSoundGate Condensor Microphone CM16/CMPA, Avisoft Bioacoustics, Berlin, Germany) suspended 30 cm above the floor of the open field arena. The procedure of USVs recording and analyses was previously described by Potasiewicz et al. (2018). In brief, the acoustic data were recorded using Avisoft RECORDER USGH (Avisoft Bioacoustics, Berlin, Germany). The spectrograms were generated with a fast Fourier transform (FFT)-length of 512 points and a time-window overlap of 75% (100% frame, Hamming window). The calls were manually marked on the computer screen and counted by an experienced user blinded to the treatment in Raven Pro Interactive Sound Analysis Software, version 1.5 (The Cornell Lab of Ornithology Bioacoustics Research Program, Ithaca, NY, USA).

The analysed USV features included a) the number of USVs, b) the average duration of the call (length of the call, measured in milliseconds), c) the bandwidth (difference between the highest and lowest frequency, a measure of frequency modulation, expressed in kHz), and d) the peak frequency (the frequency in kHz at which maximal energy occurs within the selection). Based on these features, we manually divided the USVs into two major categories: 22-kHz ‘alarms’ and 50 kHz, ‘happy’ calls. Within the 50 kHz spectrum, we further distinguished the following sub-categories: short, flat, and frequency-modulated calls: low frequency-modulated (LFM) and high frequency-modulated (HFM). A detailed description of those categories is presented in Fig. 1B. The HFM category includes trills, which are the most characteristic call type for highly-rewarding situations, such as positive social interactions (Burgdorf et al., 2008).

### Novel Object Recognition Test (65-75 PND)

The novel object recognition test was performed in the same room and apparatus as the SI test and in accordance with a previously published procedure (Ennaceur & Delacour, 1988; Potasiewicz et al., 2017). In short, the test comprised two 3-min trials separated by an inter-trial interval (ITI) of 1 h. During the first trial (familiarisation), two identical objects (A1 and A2) were presented in opposite corners of the apparatus. In the second trial (retention), one of the objects was replaced with a novel object (A = familiar and B = novel). The exploration of an object was defined as looking at, sniffing, or touching the object while sniffing but not leaning against, standing on, or sitting on the object. The data from any rat spending less than 5 s exploring the two objects during the familiarisation or retention trial was eliminated from the analysis. The behaviour was recorded by a CCTV camera placed above the arena and connected to the Any-maze® tracking system, which automatically measured the distance travelled by the tested animal. The experimenter, blinded to the treatment conditions, manually assessed the exploratory activity of animals. Based on the exploration time (E) of the two objects in the retention trial, a discrimination index (DI) was calculated as (EB– EA)/(EA+EB).

### Identification of the oestrous cycle phase

For all tested females, vaginal cytology samples were collected by the end of each experimental day to determine the stage of the standard 4-day oestrous cycle (metestrus, diestrus, proestrus, or estrus) to check whether the oestrous cycle had any influence on the social behaviour of the tested females. The procedure of identification of the oestrous cycle phase was previously described in detail by (Potasiewicz et al., 2020).

### Statistical analysis

The data on *social behaviour, ultrasonic call’ features* (duration, bandwidth, and peak frequency) and *discrimination index* in the NOR test were analysed by two-way ANOVAs with MAM and LIT-001 treatment as the between-subject factors. The *number of USVs* was analysed by a three-way ANOVA with LIT-001 and MAM treatment as between-subject factors and call category as a repeated measure.

When there was a significant main effect of treatment, we used the Tukey HSD post hoc tests to assess overall differences between treatment conditions. *A priori* planned contrast comparisons were used to compare between-group differences.

The sphericity was verified using Mauchly’s test. If the assumption of sphericity was rejected, the corrected Greenhouse–Geisser value was utilised. The effect size was estimated using partial eta squared (ŋp2).

The normality of data distribution was evaluated by the Kolmogorov-Smirnov test. Statistical significance was set at p < 0.05. The statistical analyses were performed using Statistica 12.0 for Windows. Detailed ANOVA results and the effect sizes are presented in Table S1 (Supplement). Tables containing information on the number of subjects per group in each behavioural test and on subjects excluded from the analysis are presented in Table S2 (Supplement).

## RESULTS

MAM treatment did not change the exploratory activity of rats during adaptation to the Open Field arena (Supplement, Fig. S2). Control and MAM-treated animals did not differ in weight (Supplement, Fig. S3) or physical appearance. We observed no significant effect of the oestrous cycle in any of the measured parameters, although there was a slight tendency (p = 0.0931) of LIT-001 to increase the time of social interaction selectively during the non-receptive phase (Supplement, Fig. S4).

### Social Interaction test

The data presented are a sum of all scored behaviours: sniffing, anogenital sniffing, grooming, climbing, and following, recorded during a 10-minute test session.

#### Males

As presented in Fig. 2A, the prenatal MAM treatment reduced overall time spent on social interactions, as compared to the control groups (p = 0.0269, Tukey HSD post hoc test following a significant MAM effect: F[1,93] = 4.71, p = 0.0325; Fig. 2A). LIT- 001 showed an overall trend to increase the time spent on social activities (LIT-001 effect: F[1,93] = 2.51, p = 0.0634). Planned contrast comparisons revealed that LIT- 001 at a dose of 1 mg/kg increased the time spent on social interactions in MAM animals (t = −2.52, p = 0.0131), whereas 3 mg/kg showed a tendency towards increasing the time of social activities (t = −1.91, p = 0.0580) in the control group.

**Figure 2.**
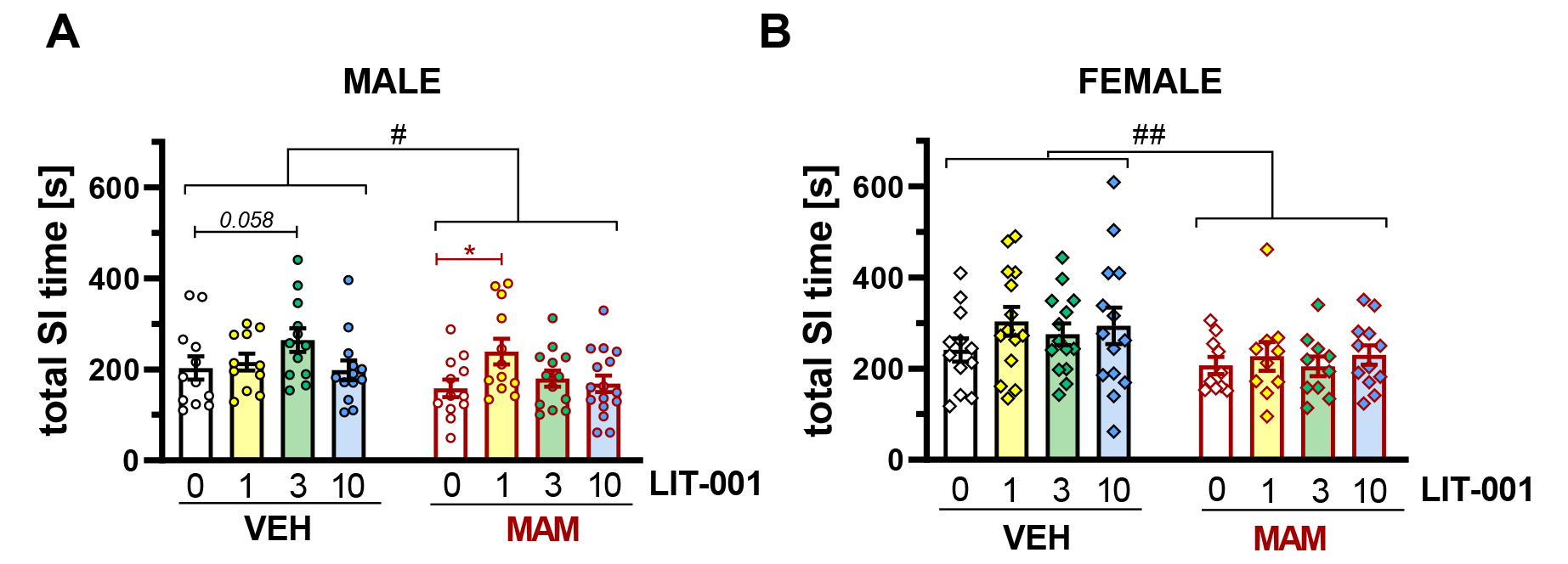
Effects of prenatal MAM exposure and treatment with oxytocin receptor antagonist LIT-001 on the total time spent on social behaviour in the social interaction test. *The total time of social activities of male (A) and female (B) animals was assessed during a 10-min session of the SI test. The calculated value is the* sum of all social interactions per pair of animals. *Bars represent the mean and S.E.M, with individual values presented as dots (males) or squares (females).* Symbols: # p < 0.05, ## p < 0.01, T*ukey* HSD; *p < 0.05, *planned contrast comparisons*.

#### Females

As shown in Fig. 2B, females presented a decrease in the time of social interactions (p = 0.0034, Tukey HSD post hoc test following a significant MAM effect: F[1,88] = 8.83, p = 0.0038; Fig. 2B). No significant effect of LIT-001 was found in this part of the study.

### Social Interaction-induced USVs

#### Males

MAM reduced the number of emitted USVs, and this reduction was significant for HFM calls (p < 0.0001, Tukey HSD post hoc test following a significant MAM x call type interactions: F[1.12, 100.11] = 4.71, p = 0.0325; Fig. 3A, Supplement, Table S3). LIT-001 administration affected USVs emission in a MAM-dependent manner (a significant LIT-001 x MAM x call type interactions: F[3.37, 100.11] = 2.87, p = 0.0343; Fig. 3A, Table). Planned contrast comparisons revealed that this effect was mainly driven by a tendency of LIT-001 at 3 mg/kg to increase vocalisations in controls (t = - 1.76, p = 0.0818).

**Figure 3.**
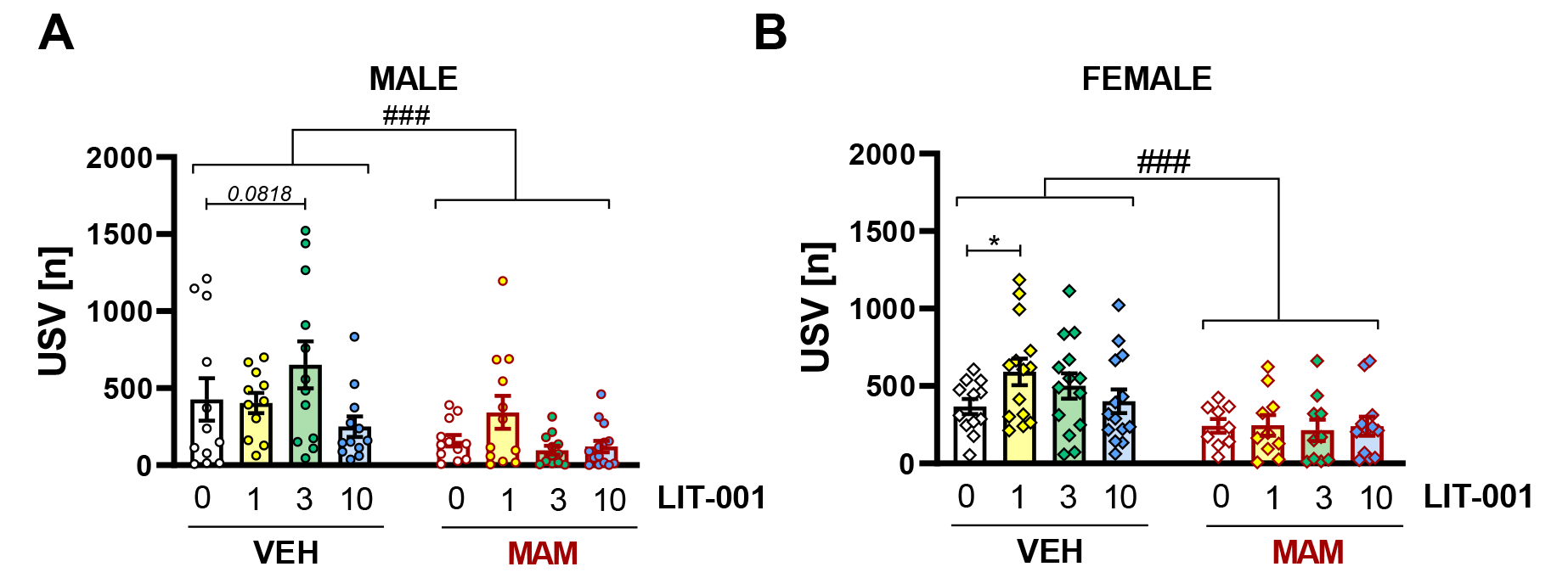
Effects of prenatal MAM exposure and LIT-001 treatment on the number of ultrasonic calls emitted during the social interaction test. *The number of high-frequency modulated calls emitted by males (A) and females (B) during a 10-minute recording of the social interaction test. Presented number of ultrasonic calls was calculated per pair of animals. Bars represent means and S.E.M, with individual values presented as dots (males) or squares (females). Symbols: ### p < 0.001, Tukey HSD; *p < 0.05, planned contrast comparisons*.

Since the effect of MAM was restricted to HFM calls, the further analysis focused on this call type. Detailed analysis of call parameters revealed a reduced bandwidth of MAM-treated males’ calls (p = 0.0003, Tukey HSD post hoc test following a significant MAM effect: F[1, 88] = 15.32, p = 0.0002; Fig. 4A). Although no significance was found, there was an overall trend of LIT-001 to increase the bandwidth of emitted USVs (F[3, 88] = 2.33, p = 0.079). Planned contrast comparisons showed a tendency of LIT-001 at 3 mg/kg to increase bandwidth (t = −1.73, p = 0.0866) in controls. No significant effects of MAM or LIT-001 were found for call duration and peak frequency (Fig. 4C and 3E).

**Figure 4.**
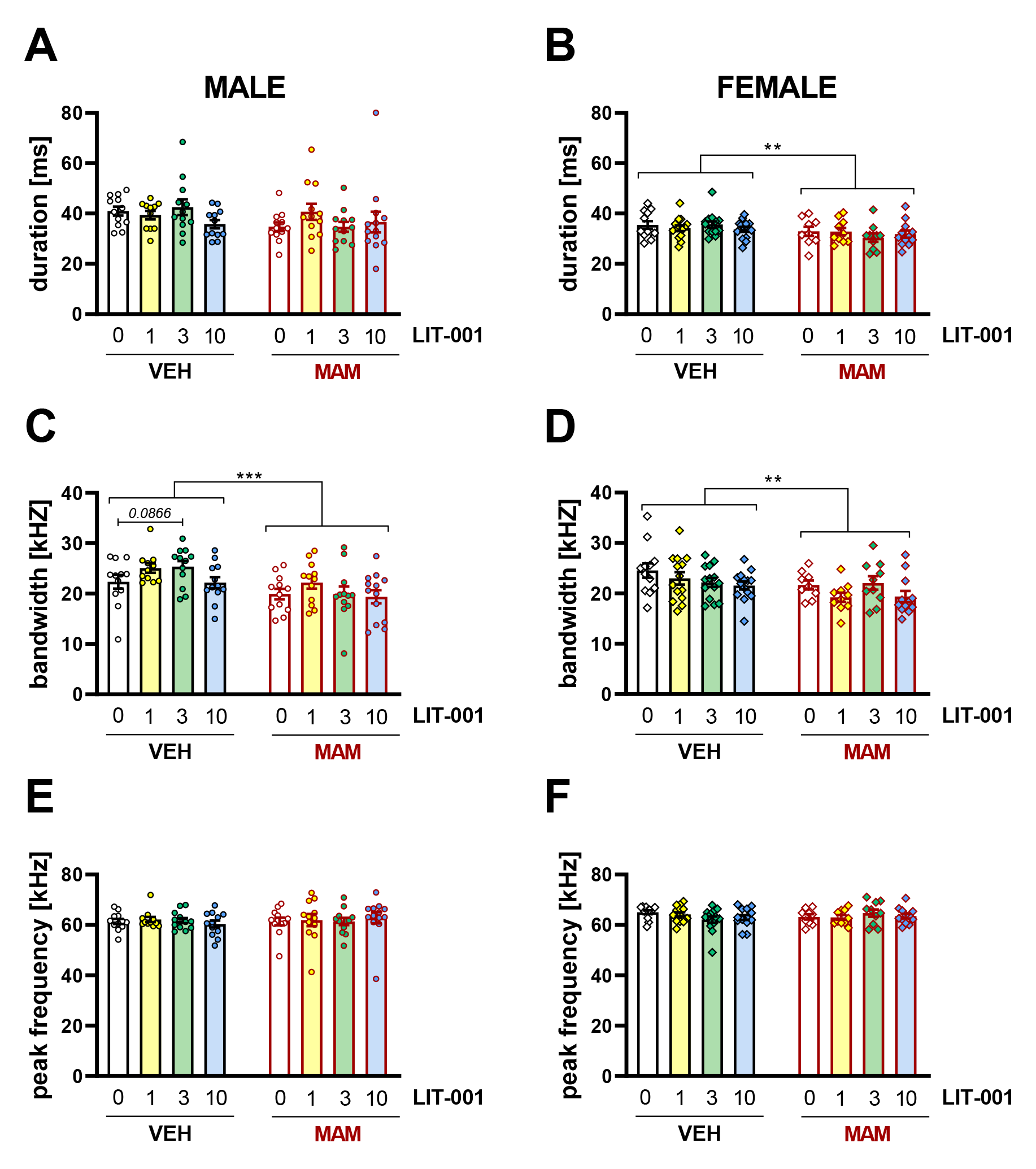
Effects of prenatal MAM exposure and LIT-001 treatment on acoustic parameters of USVs emitted during the social interaction test: mean duration, bandwidth and peak frequency. *Bandwidth (A & B), average duration (C & D), and peak frequency (E & F) of HFM calls were calculated per pair of animals during a 10-minute recording of the Social Interaction test. Results are presented separately for male (A, C & E) and female (B, D & F) pairs of animals. Bars represent means and S.E.M, with individual values presented as dots (males) or squares (females). Symbols: **p < 0.01, ***p < 0.001, Tukey HSD*.

#### Females

MAM reduced the number of USVs emitted by females, and this reduction was significant for the HFM calls (p < 0.0001, Tukey HSD post hoc test following a significant MAM x call type interactions: F[1.16, 101.54] = 18.54, p < 0.0001; Fig. 4B, Supplement, Table S3). Planned contrast comparisons revealed that LIT-001 at 1 mg/kg increased the number of HFM calls in controls (t = −2.28; p = 0.025).

The acoustic parameters of HFM calls were calculated and analysed. MAM reduced the mean duration (p = 0.0061, Tukey HSD post hoc test following a significant MAM effect: F[1, 87] = 7.85, p = 0.0063; Fig. 4D) and bandwidth (p = 0.0055, Tukey HSD post hoc test following a significant MAM effect: F[1, 87] = 7.94, p = 0.0059; Fig. 4B). No significant changes were observed for the peak frequency of emitted calls (Fig. 4F). LIT-001 treatment did not influence USVs parameters in female rats.

### Novel Object Recognition

#### Males

Based on the exploration time of the novel and familiar objects (Fig. S5C, Supplement), the discrimination index (DI) was calculated. As illustrated in Fig. 5A, MAM reduced the DI (p = 0.0001, Tukey HSD post hoc test following a significant MAM effect: F[1, 111] = 23.18, p < 0.0001; Fig. 5A). LIT-001 showed a tendency towards increasing the DI in MAM-treated males (an overall trend for LIT-001 x MAM interaction: F[3, 111] = 2.58, p = 0.057). Planned contrast comparisons revealed that LIT-001 increased the DI at 1 mg/kg (t = −2.9814, p = 0.0035) and 3 mg/kg (t = −2.73, p = 0.0072) in MAM-treated male animals.

**Figure 5.**
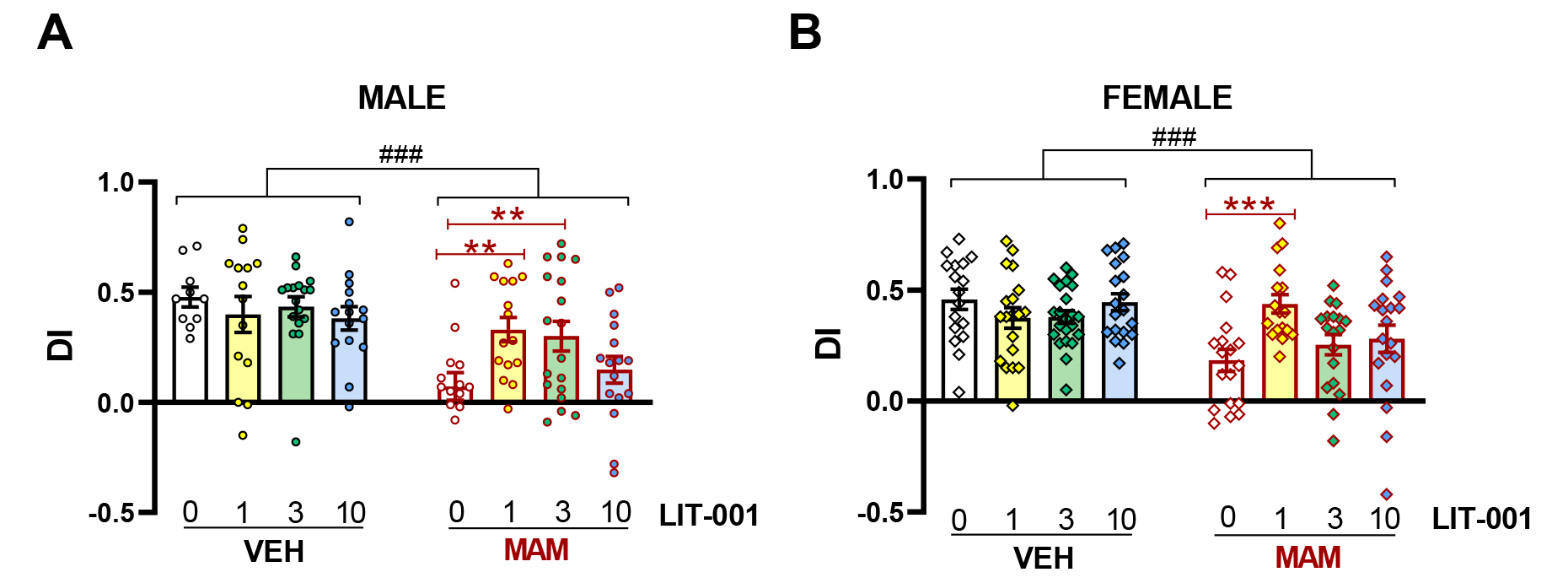
Effects of prenatal MAM exposure and LIT-001 treatment on Discrimination Index in the Novel Object Recognition test. *From the exploration times for familiar (EA) and novel (EB) objects, the Discrimination Index (DI) was calculated: DI = (EB– EA)/(EA+EB). The test stage lasted for 3 minutes. Results are presented separately for males (A) and females (B). Bars represent means and S.E.M, with individual values presented as dots (males) or squares (females). Symbols: ### p < 0.001, Tukey HSD; **p < 0.01, ***p < 0.001, planned contrast comparisons*.

#### Females

MAM reduced DI also in female rats (p = 0.0001, Tukey HSD post hoc test following a significant MAM effect: F[1, 141] = 15.18, p = 0.0002; Fig. 5B). LIT-001 treatment increased the DI in MAM-treated females (significant LIT-001 x MAM interaction: F[3, 142] = 4.63, p = 0.004). Planned comparisons revealed that this effect was specific for the dose of 1 mg/kg (t = −3.8540, p = 0.0002).

Neither of the treatments used influenced the distance travelled in the NOR test. Calculated distances for T1 and T2 are presented in Fig. S6 (Supplement). Exploration times of objects in T1 were also calculated. Neither MAM nor LIT-001 influenced exploration behaviour in T1 (Supplement, Fig. S5A, A5B).

Due to low object exploration (total exploration < 5 s or lack of exploration of one of the objects) at T2, data from 19 male animals were excluded from statistical analysis.

## DISCUSSION

The results of the present study confirm that prenatal exposure to the MAM neurotoxin induces developmental disturbances in rats resulting in social and cognitive deficits. To investigate social impairments, we used the social interaction test. MAM-treated animals spent significantly less time on social behaviours in comparison to controls, and this effect regarded both sexes. Reduction in social interaction time is further supported by the decreased amount of ‘happy’ ultrasonic calls. We also report MAM-induced deficits in cognition, assessed with the novel object recognition test. Decreased discrimination indexes were evident in the schizophrenia model regardless of sex.

A novel, nonpeptide oxytocin receptor agonist, LIT-001, was used to rescue those deficits. In the SI test, 1 mg/kg of LIT-001 increased the total time of social interactions in MAM-treated males but not in females, whereas 3 mg/kg tended to increase SI in control males. LIT-001 also affected ultrasonic vocalisations: while the dose of 1 mg/kg was effective in control females, 3 mg/kg only slightly increased some USV parameters in control males, and 10 mg/kg did not affect social behaviours or the USVs. In the cognitive part of the study, 1 mg/kg rescued cognitive impairments in DI in both sexes, while 3 mg/kg was effective only in males, and 10 mg/kg again showed no beneficial effect. Overall, dependent on the measure, we report either no dose-related effects or an inverted ‘U’ shape dose-related action.

### Social behaviour

We report decreased time spent on social interactions during the SI test by the MAM-treated animals. This effect is consistent with the previous work from our group, where MAM animals presented a reduced time of social activity both as juveniles and in adulthood (Potasiewicz et al., 2020, 2018). Furthermore, this effect was evident in both males and females. Reduction of the total time spent on social activities was not a result of decreased locomotor activity since we have observed no differences between MAM-treated and control animals in the distance travelled in the open field arena during a 5-minute adaptation on a day before the SI test (Supplement, Fig. S2). The social deficit in the MAM model was also described by (Bator et al., 2018; Flagstad et al., 2004; Le Pen et al., 2006); however, in those studies, only male animals were used. Female, but not male MAM-treated rats were tested by Hazane et al. (2009), who reported a decrease in the time spent on social interactions.

LIT-001 (1 mg/kg) increased the total time of social interactions in MAM-treated males but not in females. Interestingly, 3 mg/kg of LIT-001 slightly increased the time of social interactions in control male rats. Since, to date, this is the second work describing the effects of LIT-001 on social behaviour and the first done in rats, it is difficult to compare these results with other studies. In the original paper introducing the LIT-001 compound, a mouse genetic model of autism was used. LIT-001 acute administration restored deficits in social contacts of mice, scored in the social interaction test. The authors investigated the effects of doses 10 and 20 mg/kg of LIT-001, and the lower dose was the most effective. Both male and female animals were used, but the results were presented as a sum for both sexes; hence no information was provided on sex differences in these testing conditions (Frantz et al., 2018).

Despite the lack of other studies using the same OXTR agonist, several investigations have been made using OXT or OXT analogues. Subcutaneous OXT administration increased time spent on social interactions in the Lister-hooded rats and attenuated PCP-induced hyperlocomotion (Kohli et al., 2018). OXT administration and stimulation of OXT neurons rescued disrupted social behaviour in Cntnap2 mice, one of the genetic models of autism spectrum disorders, and the effect was blocked by a selective OXTR antagonist (Peñagarikano et al., 2015). Due to problematic Blood Brain Barrier penetration by oxytocin, most animal studies have been done using intracerebroventricular (i.c.v.) administration or, similarly to humans, nasal sprays. Acute intranasal OXT treatment increased male-female social interactions, had no effect on interactions between familiar male mice, and decreased interactions between unfamiliar males (Huang et al., 2014). Social avoidance and lack of social preference after social defeat in male rats was rescued by a single i.c.v. OXT injection (Lukas et al., 2011). Although penetration of BBB by exogenous OXT given intraperitoneally or subcutaneously is debatable, some effects of such treatments are being reported. This can be due to the peripheral action of oxytocin (for review, see Feifel et al., 2016).

### Ultrasonic communication during the Social Interaction test

During the social interaction test, the ultrasonic communication of animals was recorded. We found a significant decrease in the number of ‘happy’ calls (high-frequency modulated USVs of about 50 kHz) in both males and females treated prenatally with MAM. This finding is consistent with previous work from our group (Potasiewicz et al., 2020). Besides this report, little is known about changes in the USVs of MAM-treated animals. For other schizophrenia models, USVs were measured in pharmacological models using MK-801 and ketamine, and decreases in the number of calls were reported (Faure et al., 2019; Nikiforuk et al., 2013). In a general genetic model of neuropsychiatric diseases (Cacna1c knockout), ultrasonic communication in the 50 kHz range was diminished in juvenile rats in males but not in females (Kisko et al., 2018). Decreases in social communication were also reported in mouse genetic models of autism (Scattoni et al., 2011). A reduced number of ‘happy’ calls may be interpreted as a social communication impairment or an additional measure of social withdrawal, both of which are considered a part of social dysfunction in schizophrenia (Millan et al., 2014).

Although there was a significant interaction between MAM and LIT-001 treatment, the detailed analysis showed that this effect was mainly driven by a tendency to increase the number of calls by LIT-001 at the dose of 3 mg/kg in control male animals. No significant effect of LIT-001 in any of the tested doses of LIT-001 was seen for MAM-treated animals, neither male nor female. Interestingly, however, 1 mg/kg of LIT-001 increased the number of USVs in control females.

In addition to the number of USVs, we also looked into the specific parameters of those calls. Changes in USV structure can be interpreted as a further impairment of communication in the MAM-treated animals, which has some resemblance to the speech disruption (e.g., impoverished content of speech, blocked speech) reported in patients with schizophrenia (Messinger et al., 2011). According to our previous research, MAM-treated animals emitted shorter calls and of lower bandwidth (a measure of call complexity) than controls (Potasiewicz et al., 2020). Similarly, in the current study, MAM also decreased calls’ bandwidth in both sexes. Moreover, the mean duration of emitted USVs was decreased in MAM-treated females but not in males.

No significant effects of LIT-001 on acoustic call parameters were observed; however, there was a tendency towards increasing bandwidth in control male animals treated with 3 mg/kg of the OXTR agonist. The effects of LIT-001 on ultrasonic communication have not been investigated before. However, some findings exist on the OXT system’s influence on rodent ultrasonic calls. Acute intranasal OXT treatment did not significantly alter ultrasonic communication during social interactions between male and female mice but had a tendency towards increasing the number of calls and mean duration of emitted USVs (Huang et al., 2014). In another study, subcutaneous OXT administration, which increased time spent on social interaction, did not alter the number or specific parameters of USVs emitted during the SI test (Kohli et al., 2018).

Reports of different alterations of ultrasonic communication in OXT knockout mice suggest that the OXT system influences the USVs’ repertoire. For example, in a mother isolation test, an OXT null knockout mice pups vocalised less than wild-types and presented more aggressive behaviours as adults (Winslow et al., 2000). No sex differences in pup vocalisations were observed there. Similarly, OXTR knockout mice also emitted fewer calls during social isolation (only males were tested), and more aggressive behaviours were noted in those animals than in controls (Takayanagi et al., 2005).

In addition to the communication role, USVs analysis provides insight into animals’ emotional state, as reward-related, ‘happy’ calls have different characteristics than fear-and-aggression-related, ‘alarm’ calls. In this study, we hardly ever observed any ‘alarm’ calls, and their appearance was not correlated to any of the experimental conditions. However, the observed decrease in the number of 50 kHz calls can also be interpreted as a diminished positive valence of social interaction or loss of rewarding properties of social contexts.

### Novel Object Recognition test

In the NOR test, both male and female MAM-treated animals presented cognitive deficits. Cognitive impairments induced by prenatal MAM administration were previously reported by our group (Potasiewicz et al., 2020) and by other researchers (Bator et al., 2018; Flagstad et al., 2005).

LIT-001 improved animals’ performance in the NOR test, and 1 mg/kg had the most beneficial effect in both sexes, whereas 3 mg/kg increased DI only in males. LIT-001’s effects on animal cognition were not previously tested; however, current results are consistent with oxytocin’s effects on social recognition and memory in laboratory animals (e.g., Ferguson et al., 2001; Popik et al., 1992).

Moreover, improvement of recognition memory in NOR test, induced by an i.c.v. OXT administration was reported by Havranek et al. (2015), and intranasal oxytocin had a beneficial effect on recognition memory impairments caused by prolonged stress (Park et al., 2017) and in maternally separated rats (Joushi et al., 2021). OXT’s i.c.v. administration, as well as the OXT derivative given in a nasal spray, rescued memory impairments in a mice model of Alzheimer’s disease (Takahashi et al., 2022). Interestingly, the neuroprotective effects of OXT against cognitive dysfunction after cranial radiation were also reported (Igarashi et al., 2022).

### Doses and the inverted U hypothesis

The effects of LIT-001 reported here were not dose-dependent. We report that the most effective dose was 1 mg/kg, while 3 mg/kg showed some efficacy in males but not in females, and 10 mg/kg was ineffective. Similarly, in the mice genetic model of autism, the most significant effect was induced by a lower dose (10 vs. 20 mg/kg of LIT-001) (Frantz et al., 2018).

This dose-response can be explained by ‘the inverted U hypothesis of a sex-dependent relationship between OXT dose and its effects on social reward’ (Borland et al., 2019). According to Borland’s hypothesis, OXTs’ action in the dopamine mesolimbic system is responsible for the rewarding properties of social interactions in an inverted U, dose-response manner. Consequently, too high a dose of OXT (or OXTR agonist) will not produce a rewarding action or may even cause an aversion to social stimuli. Furthermore, this effect is sex-dependent as females are postulated to be more susceptible to oxytocin’s effects, and the rewarding doses of OXT are lower for females than for males. Borland et al. (2019) also suggested that this effect is not mediated through the vasopressin receptors, so it could apply to the LIT-001, a selective OXTR agonist.

A sex-dependent inverted U function has also been hypothesised for the relationship between brain OXT levels and neural activity in human studies (Feng et al., 2015) and was proposed for the effects on social recognition in rats, where moderate doses of OXT facilitate and high doses attenuate social recognition (Benelli et al., 1995). Similar inverted U dose-response effects of OXT were reported in zebrafish (Braida et al., 2012).

In the current study, LIT-001 in the highest dose did not induce the aversive effect, but the OXTR agonist effects on animal behaviour were non-linear and sex-dependent, which is consistent with Borland’s hypothesis. None of the chosen doses influenced the social interaction time in females; thus, we cannot exclude the possibility that a dose lower than 1 mg/kg of LIT-001 would be more effective.

## CONCLUSIONS

The current study addresses the previously reported social and cognitive deficits in the MAM model of schizophrenia being reversed with the novel OXTR agonist LIT-001, which showed some efficacy and thus could be considered as a pro-social and pro-cognitive agent. Future work could involve the use of selective OXTR agonists as additive medications to the standardised antipsychotic treatment in schizophrenia patients with social and cognitive impairments. The perspective of the positive effects of LIT-001 medication in patients with autism spectrum disorders or social phobia is also possible.

## CONFLICTS OF INTEREST

The authors declare that they have no conflicts of interest to disclose.

## ACKNOWLEDGMENTS

Funding of this study was provided by the Polish Ministry of Education and Science grant DIAMENTOWY GRANT 2019, no. 0049/DIA/2019/48 to D.P.

## Notes

### Competing Interest Statement

The authors have declared no competing interest.

