## Supplement for "Pro-social and pro-cognitive effects of LIT-001, a novel oxytocin receptor agonist in a neurodevelopmental model of schizophrenia"

**Fig. S1 Effects of MAM exposure on litter characteristics.**

Number of offspring per each dam with distinction made for male and female offspring. A two-way ANOVA revealed no effect of MAM treatment on the number of pups but a tendency towards a higher female/male ratio. Symbols: ♂ (males) and ♀ (females) represent indivual values, with a line indicating the mean value of the littersize.

| The mean number of pups (7 VEH litters and 7 MAM litters) | | |
|---|---|---|
| | VEH | MAM |
| Male | 6.57 | 7.71 |
| Female | 7.86 | 5.71 |
| **Total** | **14.29** | **13.57** |
| Male/Female ratio | 0.84 | 1.35 |

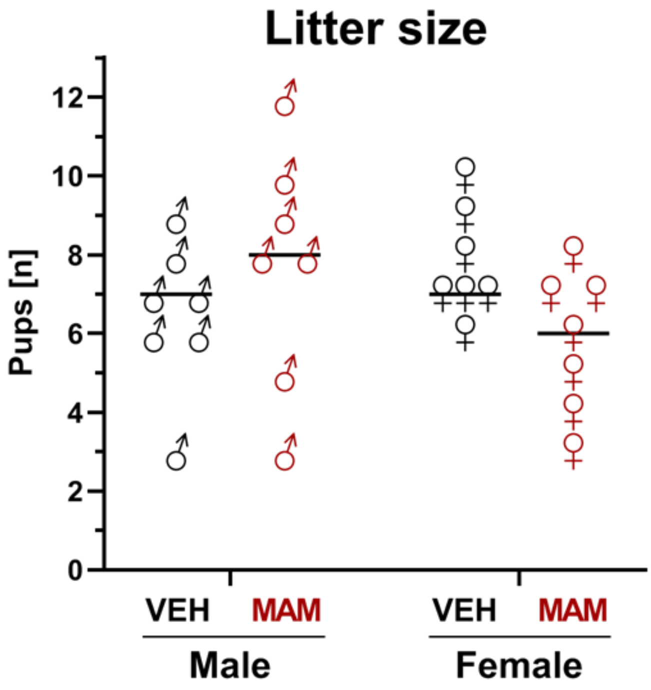


*Fig. S1. Data are presented as a mean ± SEM of the number of pups in each litter.*

| ANOVA results. | | | | | |
|---|---|---|---|---|---|
| *EFFECT* | Degr. of freedom | F | p | Partial eta-squared | Observed power |
| ***MAM treatment*** | 1, 24 | 0,1995 | 0,6592 | 0,0082 | 0,0713 |
| ***Sex*** | 1, 24 | 0,3910 | 0,5377 | 0,0160 | 0,0922 |
| ***MAM*Sex*** | 1, 24 | 4,2207 | 0,0510 | 0,1496 | 0,5048 |

**Table S1 Results for statistical analysis for USVs characteristics.**

Data were analyzed by two-way ANOVAs with the MAM treatment (VEH and MAM) and LIT-001 treatment (0, 1, 3, and 10 mg/kg) as the between-subject factors. For the USVs analysis, three-way ANOVA was used with the MAM treatment (VEH and MAM), LIT-001 treatment (0, 1, 3, and 10 mg/kg), and call category (short, flat, low frequency-modulated, and high frequency-modulated) as the between-subject factors.

1. **Social behavior during the social interaction test**

**Males**

| *EFFECT* | Degr. of freedom | F | p | Partial eta-squared | Observed power |
|---|---|---|---|---|---|
| ***MAM treatment*** | 1, 93 | 4.7105 | 0.0325 | 0.0482 | 0.5746 |
| ***LIT-001 treatment*** | 1, 93 | 2.5117 | 0.0634 | 0.0750 | 0.6048 |
| ***MAM*LIT*** | 3, 93 | 1.9201 | 0.1317 | 0.0583 | 0.4818 |

**Females**

| *EFFECT* | Degr. of freedom | F | p | Partial eta-squared | Observed power |
|---|---|---|---|---|---|
| ***MAM treatment*** | 1, 88 | 8.8284 | 0.0038 | 0.0912 | 0.8361 |
| ***LIT-001 treatment*** | 3, 88 | 0.8666 | 0.4616 | 0.0287 | 0.2320 |
| ***MAM*LIT*** | 3, 88 | 0.1984 | 0.8972 | 0.0067 | 0.0857 |

2. **USVs emitted during the social interaction test**

a. **The number of 50 kHz calls**

**Males**

| *EFFECT* | Degr. of freedom | F | p | Partial eta-squared | Observed power |
|---|---|---|---|---|---|
| ***MAM treatment*** | 1, 89 | 14.6166 | 0.0002 | 0.1411 | 0.9657 |
| ***LIT-001 treatment*** | 3, 89 | 1.7500 | 0.1621 | 0.0557 | 0.4425 |
| *Calltype* | 3, 267 | 81.1605 | 0.0000 | 0.4770 | 1.0000 |
| ***MAM*LIT*** | 3, 89 | 3.0107 | 3.0343 | 0.0921 | 0.6918 |
| ***MAM*Calltype*** | 3, 267 | 14.9679 | 0.0000 | 0.1440 | 1.0000 |
| ***LIT*Calltype*** | 9, 267 | 2.0906 | 0.0306 | 0.0658 | 0.8683 |
| ***MAM*LIT*Calltype*** | 9, 267 | 2.8727 | 0.0030 | 0.0883 | 0.9624 |

**Females**

| *EFFECT* | Degr. of freedom | F | p | Partial eta-squared | Observed power |
|---|---|---|---|---|---|
| *MAM treatment* | 1, 87 | 17.4741 | 0.0001 | 0.1673 | 0.9851 |
| *LIT-001 treatment* | 3, 87 | 0.9284 | 0.4306 | 0.0310 | 0.2466 |
| *Calltype* | 3, 261 | 150.9441 | 0.0000 | 0.6344 | 1.0000 |
| *MAM*LIT* | 3, 87 | 0.9983 | 0.3976 | 0.0333 | 0.2633 |
| *MAM*Calltype* | 3, 261 | 18.5430 | 0.0000 | 0.1757 | 1.0000 |
| *LIT*Calltype* | 9, 261 | 0.8973 | 0.5282 | 0.0300 | 0.4443 |
| *MAM*LIT*Calltype* | 9, 261 | 0.9304 | 0.4991 | 0.0311 | 0.4607 |

**b. The acoustic characteristics of emitted calls**

**i) Duration**

**Males**

| *EFFECT* | Degr. of freedom | F | p | Partial eta-squared | Observed power |
|---|---|---|---|---|---|
| *MAM treatment* | 1, 88 | 2.5793 | 0.1118 | 0.0285 | 0.3553 |
| *LIT-001 treatment* | 3, 88 | 0.7462 | 0.5274 | 0.0248 | 0.2037 |
| *MAM*LIT* | 3, 88 | 1.6179 | 0.1910 | 0.0523 | 0.4117 |

**Females**

| *EFFECT* | Degr. of freedom | F | p | Partial eta-squared | Observed power |
|---|---|---|---|---|---|
| *MAM treatment* | 1, 87 | 7.8520 | 0.0063 | 0.0828 | 0.7914 |
| *LIT-001 treatment* | 3, 87 | 0.3931 | 0.7583 | 0.0134 | 0.1250 |
| *MAM*LIT* | 3, 87 | 0.7646 | 0.5169 | 0.0257 | 0.2079 |

**ii) Bandwidth**

**Males**

| *EFFECT* | Degr. of freedom | F | p | Partial eta-squared | Observed power |
|---|---|---|---|---|---|
| *MAM treatment* | 1, 88 | 15.3175 | 0.0008 | 0.1483 | 0.9720 |
| *LIT-001 treatment* | 3, 88 | 2.3338 | 0.0794 | 0.0737 | 0.5687 |
| *MAM*LIT* | 3, 88 | 0.6524 | 0.5836 | 0.0218 | 0.1821 |

**Females**

| *EFFECT* | Degr. of freedom | F | p | Partial eta-squared | Observed power |
|---|---|---|---|---|---|
| *MAM treatment* | 1, 87 | 7.9396 | 0.0060 | 0.0836 | 0.7958 |
| *LIT-001 treatment* | 3, 87 | 2.1214 | 0.1033 | 0.0682 | 0.5244 |
| *MAM*LIT* | 3, 87 | 0.9774 | 0.4072 | 0.0326 | 0.2583 |

#### iii) Peak Frequency

**Males**

| *EFFECT* | Degr. of freedom | F | p | Partial eta-squared | Observed power |
|---|---|---|---|---|---|
| *MAM treatment* | 1, 88 | 0.1019 | 0.7503 | 0.0012 | 0.0615 |
| *LIT-001 treatment* | 3, 88 | 0.0751 | 0.9733 | 0.0026 | 0.0630 |
| *MAM*LIT* | 3, 88 | 0.4105 | 0.7458 | 0.0138 | 0.1287 |

**Females**

| *EFFECT* | Degr. of freedom | F | p | Partial eta-squared | Observed power |
|---|---|---|---|---|---|
| *MAM treatment* | 1, 87 | 0.0005 | 0.9817 | 0.00001 | 0.0501 |
| *LIT-001 treatment* | 3, 87 | 0.2553 | 0.8573 | 0.0087 | 0.0968 |
| *MAM*LIT* | 3, 87 | 1.7213 | 0.1685 | 0.0560 | 0.4355 |

### 3. Discrimination Index calculated during T2 in the Novel Object Recognition task

**Males**

| *EFFECT* | Degr. of freedom | F | p | Partial eta-squared | Observed power |
|---|---|---|---|---|---|
| *MAM treatment* | 1, 87 | 23.1823 | 0.000005 | 0.1728 | 0.9975 |
| *LIT-001 treatment* | 3, 87 | 1.6810 | 0.1752 | 0.0435 | 0.4300 |
| *MAM*LIT* | 3, 87 | 2.5775 | 0.0573 | 0.0651 | 0.6205 |

**Females**

| *EFFECT* | Degr. of freedom | F | p | Partial eta-squared | Observed power |
|---|---|---|---|---|---|
| *MAM treatment* | 1, 87 | 15.1806 | 0.0002 | 0.0972 | 0.9719 |
| *LIT-001 treatment* | 3, 87 | 1.6806 | 0.1739 | 0.0345 | 0.4328 |
| *MAM*LIT* | 3, 87 | 4.6301 | 0.0040 | 0.0897 | 0.8843 |

**Table S2 Number of subjects in each experimental group.**

Number of subjects (or pairs) per each experimental group and test, with information on subjects excluded from statistical analysis (outliers).

1. SI behavior Males

| Males | VEH | | | | MAM | | | |
|---|---|---|---|---|---|---|---|---|
| Treatment | 0 | 1 | 3 | 10 | 0 | 1 | 3 | 10 |
| N: | 12 | 11 | 12 | 13 | 13 | 13 | 14 | 16 |
| Excluded: | - | - | - | - | 1 | 1 | 1 | - |
| **Final N:** | **12** | **11** | **12** | **13** | **12** | **12** | **13** | **16** |

2. SI behavior Females

| Females | VEH | | | | MAM | | | |
|---|---|---|---|---|---|---|---|---|
| Treatment | 0 | 1 | 3 | 10 | 0 | 1 | 3 | 10 |
| N: | 12 | 14 | 14 | 14 | 10 | 10 | 10 | 12 |
| Excluded: | - | - | - | - | - | - | - | - |
| Final N: | **12** | **14** | **14** | **14** | **10** | **10** | **10** | **12** |

3. SI USV Males

| Males | VEH | | | | MAM | | | |
|---|---|---|---|---|---|---|---|---|
| Treatment | 0 | 1 | 3 | 10 | 0 | 1 | 3 | 10 |
| N: | 12 | 11 | 12 | 13 | 13 | 13 | 14 | 16 |
| Excluded: | - | - | - | 1 | 1 | 1 | 2 | 2 |
| Final N: | **12** | **11** | **12** | **12** | **12** | **12** | **12** | **14** |

4. SI USV Females

| Males | VEH | | | | MAM | | | |
|---|---|---|---|---|---|---|---|---|
| Treatment | 0 | 1 | 3 | 10 | 0 | 1 | 3 | 10 |
| N: | 12 | 14 | 14 | 14 | 10 | 10 | 10 | 12 |
| Excluded: | - | - | - | - | 1 | - | - | - |
| Final N: | **12** | **14** | **14** | **14** | **9** | **10** | **10** | **12** |

5. NOR Males

| Males | VEH | | | | MAM | | | |
|---|---|---|---|---|---|---|---|---|
| Treatment | 0 | 1 | 3 | 10 | 0 | 1 | 3 | 10 |
| N: | 10 | 14 | 19 | 15 | 14 | 15 | 19 | 16 |
| Excluded: | - | - | 2 | - | - | - | 1 | - |
| Final N: | **10** | **14** | **17** | **15** | **14** | **15** | **18** | **16** |

6. NOR Females

| Males | VEH | | | | MAM | | | |
|---|---|---|---|---|---|---|---|---|
| Treatment | 0 | 1 | 3 | 10 | 0 | 1 | 3 | 10 |
| N: | 17 | 19 | 22 | 19 | 18 | 18 | 18 | 19 |
| Excluded: | - | - | - | - | - | 1 | - | - |
| Final N: | **17** | **19** | **22** | **19** | **18** | **17** | **18** | **19** |

**Fig. S2 Effects of prenatal MAM exposure on exploratory activity.**

The exploratory activity of animals, measured as distance traveled in the Open Field arena, was assessed during 5-minute adaptation to the apparatus on a day before the first SI test. MAM treatment did not influence the exploratory activity regardless of sex. Bars represent means and S.E.M, with individual values presented as dots (males) or squares (females). Symbols: ***p < 0.001 vs. VEH controls (two-way ANOVA for MAM treatment and sex effects).

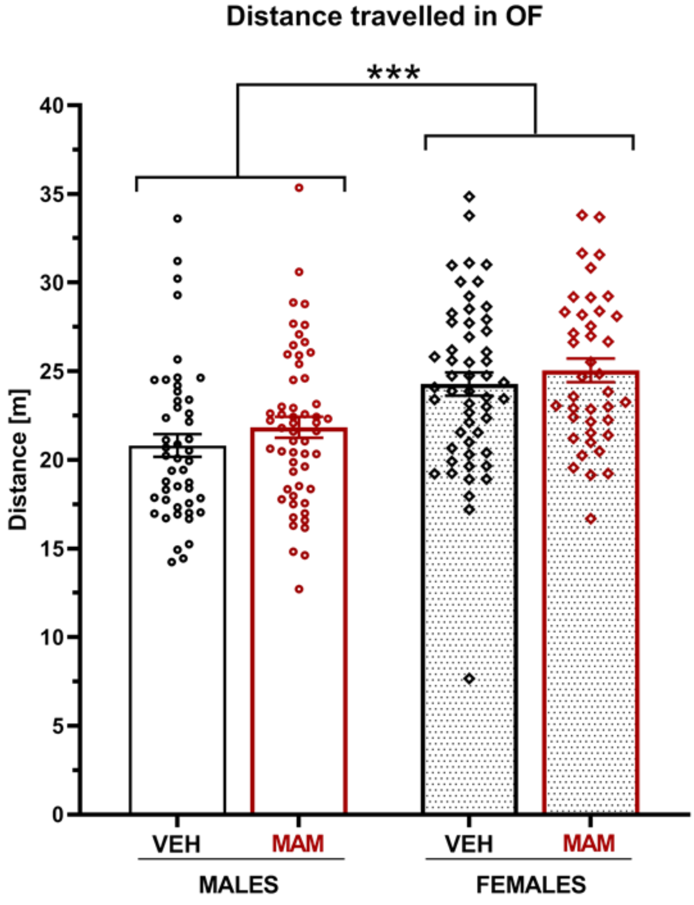


*Fig. S2.Data are presented as a mean ± SEM of the distance traveled by rats in the open field.*

| ANOVA results. | | | | | |
|---|---|---|---|---|---|
| *EFFECT* | Degr. of freedom | F | p | Partial eta-squared | Observed power |
| *Sex* | 1, 191 | 26,9897 | 0,0000 | 0,1238 | 0,9993 |
| *MAM treatment* | 1, 191 | 1,9590 | 0,1632 | 0,0102 | 0,2856 |
| *MAM*Sex* | 1, 191 | 0,0406 | 0,8404 | 0,0002 | 0,0546 |

**Fig. S3 Effects of prenatal MAM exposure on animal weights.**

The weight of all animals was measured on the day of the first SI test. MAM treatment did not influence weight of animals. The box-and-whiskers plot represent means, S.E.M and range (whiskers). Symbols: ****p < 0.00001 for the two-way ANOVA for MAM treatment and sex effects.

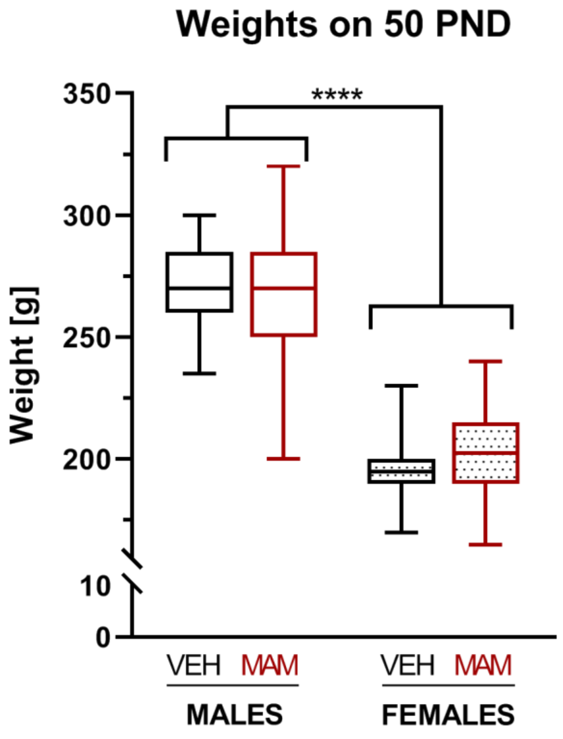


*Fig. S3. Animals weights on the day of SI test. Data presented as mean ± SEM.*

| ANOVA results. | | | | | |
|---|---|---|---|---|---|
| ***EFFECT*** | **Degr. of freedom** | **F** | **p** | **Partial eta-squared** | **Observed power** |
| ***Sex*** | 1, 191 | 566,0017 | 0,0000 | 0,7477 | 1,0000 |
| ***MAM treatment*** | 1, 191 | 0,0028 | 0,9581 | 0,0000 | 0,0503 |
| ***MAM*Sex*** | 1, 191 | 3,2517 | 0,0729 | 0,0167 | 0,4343 |

**Fig. S4 Effects of the estrous cycle on the total time of social interactions.**

The social interaction time per single animal scored during the 10-min session of the Social Interaction Test. The females were divided into two groups based on their estrous cycle phase (vaginal smears taken at the end of experimental day). The three-way ANOVA revealed a significant prenatal treatment effect and a tendency for LIT-001 (1 mg/kg) to work more effectively in females being in the "receptive" cycle phase. Bars represent means and S.E.M, with individual values presented as squares.

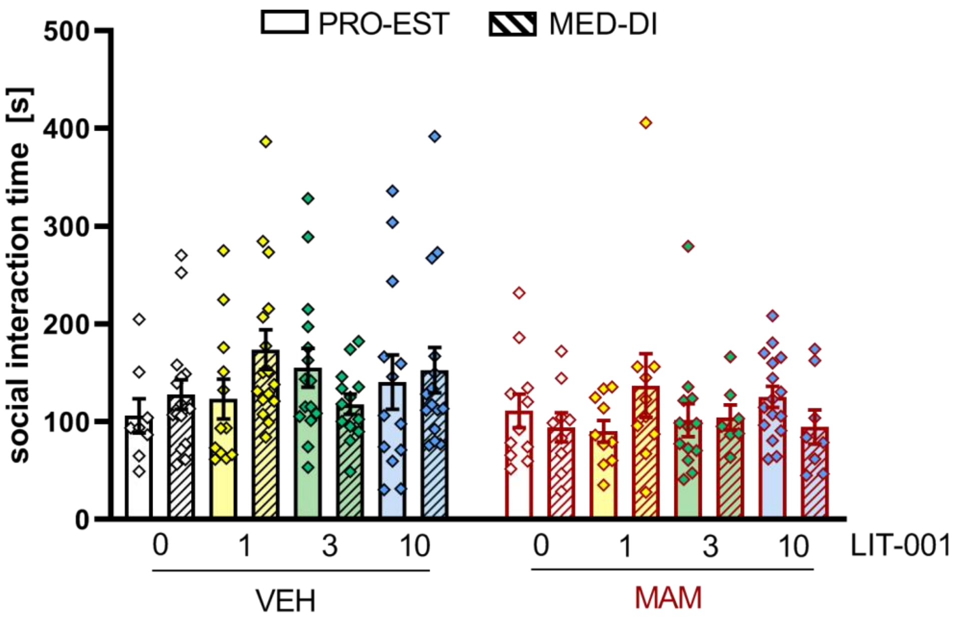


*Fig. S4. Data are presented as a mean ± SEM for the social interaction per single animal in the Social Interaction test.*

| ANOVA results. | | | | | |
|---|---|---|---|---|---|
| ***EFFECT*** | **Degr. of freedom** | **F** | **p** | **Partial eta-squared** | **Observed power** |
| ***MAM treatment*** | 1, 176 | 8.6610 | 0.0037 | 0.0469 | 0.8332 |
| ***LIT-001 treatment*** | 3, 176 | 0.8613 | 0.4623 | 0.0145 | 0.2353 |
| *Cycle* | 1, 176 | 0.3690 | 0.5443 | 0.0021 | 0.0928 |
| ***MAM*LIT*** | 3, 176 | 0.2519 | 0.8599 | 0.0043 | 0.0973 |
| ***MAM*Cycle*** | 1, 176 | 0.3168 | 0.5743 | 0.0018 | 0.0866 |
| ***LIT*Cycle*** | 3, 176 | 2.1716 | 0.0931 | 0.0357 | 0.5454 |
| ***MAM*LIT*Cycle*** | 3, 176 | 0.8971 | 0.4439 | 0.0151 | 0.2440 |

**Table S3 Number of USVs per category.**

A table containing detailed number of ultrasonic calls per each category and experimental group, emitted during the Social Interaction test.

| Males | | | Number of 50 kHz calls per category (Mean ± S.E.M) | | | | |
|---|---|---|---|---|---|---|---|
| Group | LIT-001 | N | Short | Flat | LFM | **HFM** | Total |
| VEH | 0 mg/kg | 12 | 61.92 ± 14.25 | 28.08 ± 11.94 | 135.92 ± 54.92 | **426.08 ± 137.68** | 652.00 ± 212.18 |
| | 1 mg/kg | 11 | 79.00 ± 12.16 | 4.91 ± 1.84 | 52.09 ± 12.71 | **404.09 ± 65.95** | 540.09 ± 76.35 |
| | 3 mg/kg | 12 | 95.08 ± 18.77 | 13.58 ± 4.39 | 103.50 ± 23.60 | **650.92 ± 152.72** | 863.08 ± 180.92 |
| | 10 mg/kg | 12 | 56.00 ± 9.75 | 11.17 ± 2.34 | 46.67 ± 15.08 | **250.17 ± 66.76** | 364.00 ± 84.16 |
| MAM | 0 mg/kg | 12 | 51.67 ± 9.88 | 4.00 ± 1.64 | 28.08 ± 11.03 | **158.83 ± 36.98** | 242.58 ± 51.85 |
| | 1 mg/kg | 12 | 65.92 ± 23.78 | 8.00 ± 2.20 | 97.50 ± 31.19 | **342.67 ± 107.44** | 514.08 ± 151.61 |
| | 3 mg/kg | 12 | 34.58 ± 8.21 | 3.17 ± 0.88 | 13.33 ± 4.59 | **97.42 ± 28.54** | 148.50 ± 38.58 |
| | 10 mg/kg | 14 | 43.57 ± 11.29 | 5.36 ± 1.82 | 24.93 ± 9.76 | **121.14 ± 37.24** | 195.00 ± 55.93 |
| **Females** | | | **Number of 50 kHz calls per category (Mean ± S.E.M)** | | | | |
| Group | LIT-001 | N | Short | Flat | LFM | **HFM** | Total |
| VEH | 0 mg/kg | 12 | 95.67 ± 11.86 | 13.75 ± 3.54 | 61.83 ± 14.64 | **366.83 ± 48.47** | 538.08 ± 64.80 |
| | 1 mg/kg | 14 | 160.93 ± 21.20 | 7.29 ± 1.35 | 77.00 ± 21.39 | **591.29 ± 86.56** | 836.50 ± 118.90 |
| | 3 mg/kg | 14 | 128.43 ± 17.12 | 14.43 ± 3.73 | 74.07 ± 14.16 | **500.07 ± 82.29** | 717.00 ± 98.16 |
| | 10 mg/kg | 14 | 111.64 ± 18.13 | 8.64 ± 2.05 | 59.64 ± 11.42 | **401.86 ± 76.51** | 581.79 ± 99.62 |
| MAM | 0 mg/kg | 9 | 77.89 ± 11.60 | 10.00 ± 2.93 | 45.33 ± 12.10 | **243.11 ± 44.54** | 376.33 ± 64.17 |
| | 1 mg/kg | 10 | 79.80 ± 13.02 | 8.00 ± 1.81 | 54.50 ± 16.63 | **246.40 ± 66.49** | 388.70 ± 92.67 |
| | 3 mg/kg | 10 | 73.20 ± 23.91 | 7.90 ± 2.27 | 58.10 ± 24.27 | **214.00 ± 69.14** | 353.20 ± 106.35 |
| | 10 mg/kg | 12 | 85.58 ± 12.21 | 14.92 ± 3.25 | 55.00 ± 17.77 | **241.00 ± 61.78** | 396.50 ± 81.81 |

*Table S3. Number of calls per category. Data presented as a mean ± SEM per a pair of animals in the Social Interaction test.*

**Fig S5. Effects of MAM prenatal exposure and treatment with LIT-001 on exploration times of each object during both stages of the Novel Object Recognition task.**

Exploration times for objects in the acclimatization (T1) and test stage (T2) were calculated. There was no effect of treatment (MAM or LIT-001) on the exploration time in this stage of the NOR task. No preference towards any of the objects was visible. In T2 animals spent more time exploring the novel object (hatched bars). Bars represent means and S.E.M, with individual values presented as dots (males) or squares (females). Symbols: *p < 0.05, **p < 0.01, ***p < 0.001 vs. familiar object (a paired T-test).

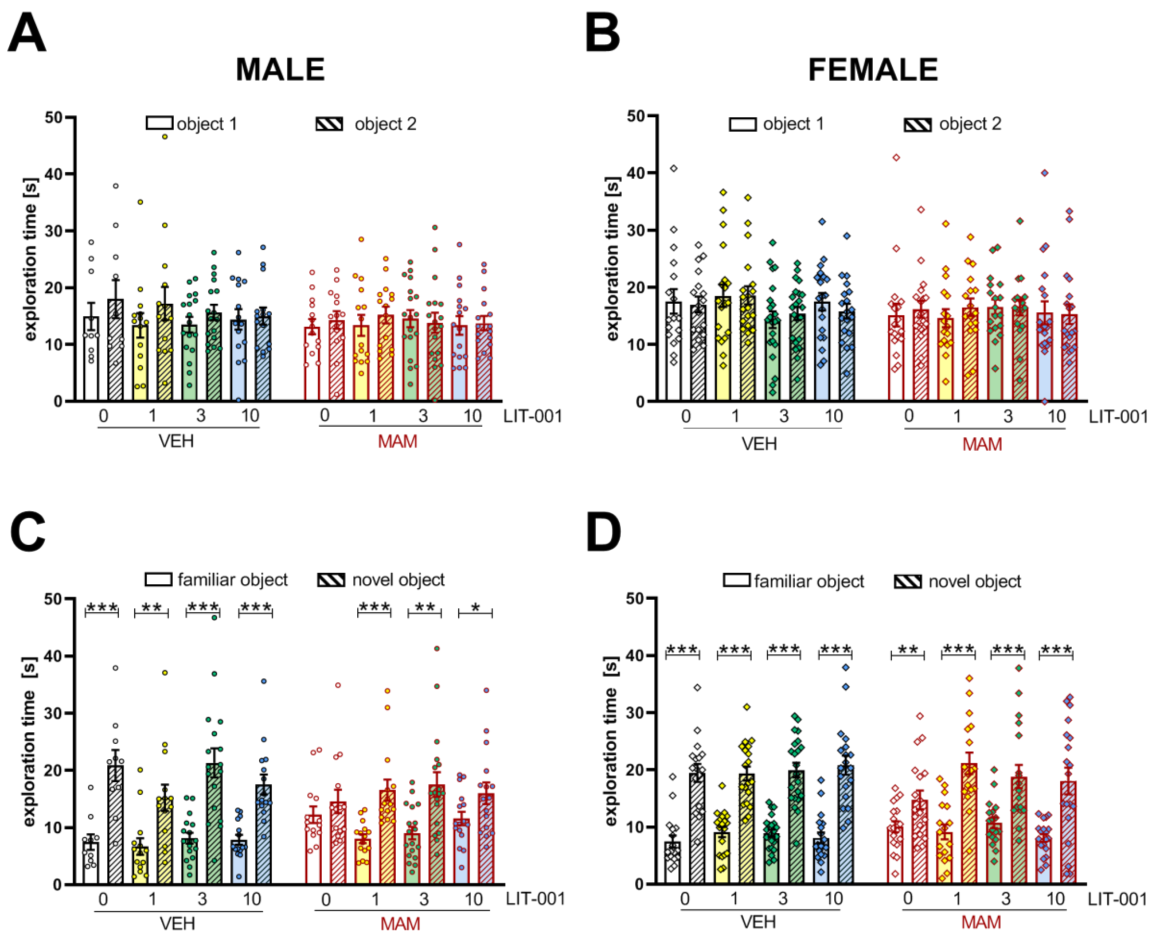


*Fig. S5. Data are presented as a mean ± SEM. Symbols: ***p<0.001, **p<0.01, *p<0.05.*

**Fig. S6 Effects of MAM prenatal exposure and treatment with LIT-001 on distance traveled by animals during both stages of the Novel Object Recognition task.**

Distance traveled by animals was calculated automatically for both stages of the NOR task: the acclimatization stage (T1; A & B) and the test stage (T2; C & D). Results are presented for both males (A, C) and females (B, D). Neither MAM nor LIT-001 treatments influenced the distance traveled by animals in the NOR task. Bars represent means and S.E.M, with individual values presented as dots (males) or squares (females).

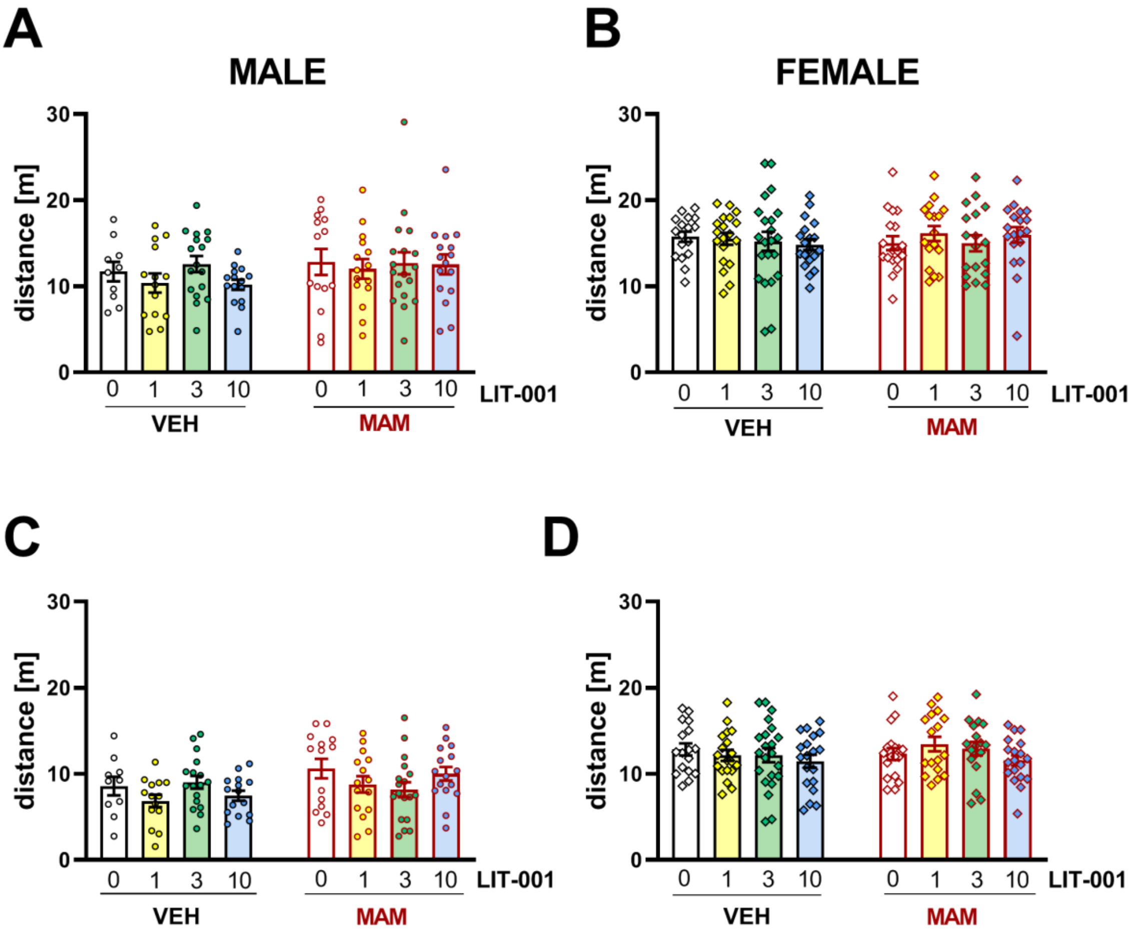


*Fig. S6. Data are presented as a mean ± SEM.*
